# Probing visual sensitivity and attention in mice using reverse correlation

**DOI:** 10.1101/2022.09.08.507101

**Authors:** Jonas Lehnert, Kuwook Cha, Kerry Yang, Daniel F. Zheng, Anmar Khadra, Erik P. Cook, Arjun Krishnaswamy

## Abstract

Visual attention is a fundamental cognitive operation that allows the brain to evoke behaviors based on the most important stimulus features. Although mouse models offer immense potential to gain a circuit-level understanding of this phenomenon, links between visual attention and behavioral decisions in mice are not well understood. Here, we describe a new behavioral task for mice that addresses this limitation. We trained mice to detect weak vertical bars in a background of checkerboard noise while audiovisual cues manipulated their spatial attention. We then modified a reverse correlation method from human studies to link behavioral decisions to stimulus locations and features. We show that mice attended to stimulus locations just rostral of their optical axis, which was highly sensitive for vertically oriented stimulus energy whose spatial frequency matched those of the weak vertical bars. We found that the tuning of sensitivity to orientation and spatial frequency grew stronger during training, was multiplicatively scaled with attention, and approached that of an ideal observer. These results provide a new task to measure spatial- and feature-based attention in mice which can be leveraged with new recording methods to uncover attentional circuits.

## Introduction

Attention is thought to be the fundamental cognitive operation that allows the brain to evoke behaviors based on the most important visual features^1–3^(**Fig 1A**). Much of what we know about the neural mechanisms of attention comes from pioneering electrophysiological work in primates which studied the effects of cueing on visually guided behavior and single neuron responses in visual cortical areas^1,2,4,5^. A common view of this work portrays attention as a spotlight that enhances neural signals about visual locations or features that are relevant to the task at hand^1,3^. This influential perspective inspired detailed models^1,6–9^ of attention and led to discoveries that attention’s effects grow as one moves from lower to higher visual areas^1,10^. However, a circuit-level, mechanistic understanding of how attention achieves its effects is still not clear.

**Figure 1.**
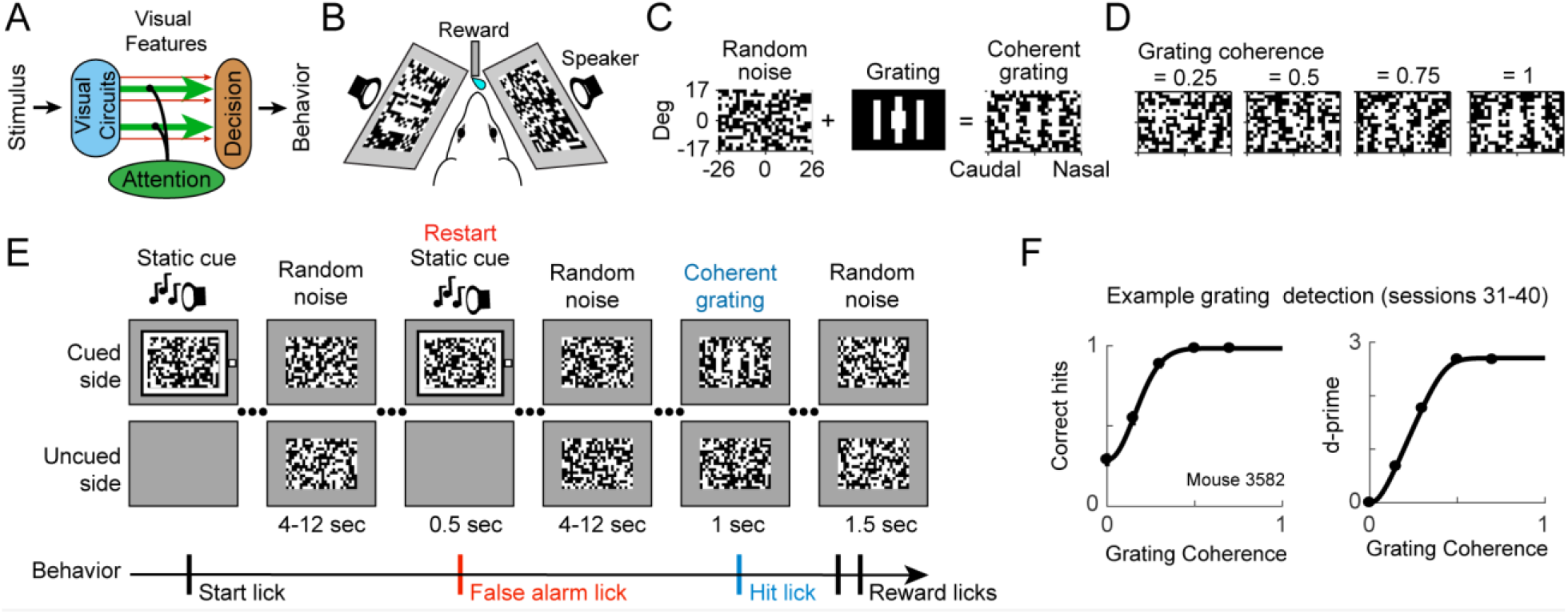
Behavioral task to measure attention in mice. **A**. Schematic shows how visual attention enhances specific visual features to aid decision making and direct behavior. **B**. Head-fixed mice faced angled visual screens (100 by 64 deg of visual angle). Speakers were located behind the screens. **C**. Dynamic checkerboard noise was combined with a 3-bar vertical grating to create the coherent grating stimulus. Checkerboard size in degrees of visual angle is indicated (optical axes = 0,0) and examples show gratings of different coherences. **D**. Schematic of our behavioral task. Trials begin with a static checkerboard and audiovisual cue that indicated the eventual location of the coherent stimulus (valid cue). In some trials the cue was invalid 10% of the time. The cue faded after mice licked (start lick) and was replaced by dynamic checkerboard noise (30 Hz stimulus update on each screen) that was presented for a randomly chosen wait period (flat hazard function). The cue was restarted if mice licked during the wait period (false alarm lick). A lick following coherent stimulus presentation (hit lick) resulted in a liquid reward (reward licks). Cue and coherent stimulus switched screens after ∼25 completed trials. **F**. Typical psychometric curves for a single well-trained mouse for sessions 31 to 40. Hits and d-prime versus stimulus coherence for valid cues (SEMs are smaller than points).

New advances in mice^11–15^, such as the ability to mark, monitor and perturb the neural signals of thousands of neurons in awake behaving animals could offer a route to make progress. Our understanding of how rodents allocate attentional resources to shape behavior, however, is still limited. A set of recent studies suggest that mice can allocate voluntary (endogenous) attention in response to a cue and can switch its spatial location between visual hemifields^16–25^. Simultaneous behavioral measurements and neural recordings in mice^16,20,23^ are beginning to identify brain regions involved in attention and resemble some findings from the classical primate literature^1^.

Current behavioral tasks, such as the Posner valid/invalid cueing paradigm^26^, usually do not simultaneously monitor a broad range of visual features to identify the ones most affected by attention (**Fig 1A**). Here, we address this issue by modifying a behavioral reverse-correlation approach that has been successfully used in humans^7,27–33^. This approach let us link multiple visual locations and features to visually guided behavior.

We trained mice to detect weak vertical bars in a background of random checkerboard noise and used audiovisual cues to manipulate their spatial attention. We then modified a reverse correlation method from human studies to relate mouse behavioral decisions to stimulus location and visual features. This tool revealed that mice did not attend to the full noise stimulus but limited their behavioral sensitivity to an ∼30 deg patch just rostral of their optical axis. In this region, sensitivity peaked for vertically oriented noise patterns whose spatial frequency matched that of the weak vertical bars. This sensitivity grew over training, predicted detection performance, and followed the location of the cue. The tuning of mouse sensitivity to stimulus features approached that of an ideal observer, and was remarkably reminiscent of human sensitivity, and demonstrated multiplicative scaling with spatial attention.

## Results

### Mice performed a spatially-cued detection task

We describe a method to reverse correlate mouse sensitivity to different features in a visual stimulus. Mice were trained to perform a cued detection task while viewing two random checkerboard stimuli (52 × 34 deg centered on the optical axis) presented on gray backgrounds (**Fig. 1B**). Briefly, animals licked a reward spout when they detected a coherent grating that appeared in one of the random checkerboard patterns. The coherent grating was produced by combining random checkerboard noise with a 3-bar vertical grating to create a range of grating stimulus strengths (referred to as grating coherence, **Fig. 1C**).

Trials began with a static checkerboard plus a visual and auditory cue on one monitor (**Fig. 1D**). This spatial cue indicated which checkerboard would eventually contain the coherent grating. The cue was always valid for the first half of the training sessions. For the second half of training sessions, we also measured attentional effects using the Posner valid/invalid cueing paradigm (the coherent grating appeared on the uncued side 10% of the time). Mice initiated the dynamic checkerboard noise by licking the reward spout which also caused the audiovisual cue to fade over 6 secs (*start lick*). If the mouse licked before the onset of the coherent grating (false alarm or *FA lick*), the single static checkerboard and audiovisual cue were re-displayed, and the trial restarted. Trials kept restarting until no FAs licks occurred. The coherent grating appeared at a random time within 4 to 12 seconds (derived from a flat hazard function). The coherent grating had a duration of one second and the mouse was rewarded for licking within a reaction time window of 0.2 to 1.5 secs (*hit lick*). No reward was given if the mouse failed to lick in response to the coherent grating (*miss*). Cues remained on the same screen for a block of ∼25 completed trials before switching to the other screen.

As shown by the example psychometric curves in **Fig. 1E**, this was a challenging detection task. We always included a grating coherence = 0 to measure the effect of FAs on the hit rate, which allowed us to report d-prime performance levels. It usually took about 20 training sessions for a typical mouse to detect a grating coherence = 0.3 at a d-prime above one.

### Using reverse-correlation to measure sensitivity and attention in mice

On average, mice produced false alarms at a rate of 0.18 FA licks per second of dynamic checkerboard noise (standard deviation = 0.04, N = 13 mice). We wondered if these FA licks occurred because the mice thought that the random noise resembled the 3-bar vertical grating they were trained to detect. If true, what features/locations of the checkerboard noise were they monitoring? To answer this, we modified a reverse-correlation technique designed to measure behavioral sensitivity and attention in humans^28,31–33^. Reverse correlation links a subject’s response to the energy contained in the preceding noisy stimulus.

Our dynamic checkerboard noise was an ideal stimulus for reverse correlation approach because it contained all spatial frequencies, orientations, and contrast patterns, and thus was statistically unbiased. Another benefit of using random checkerboard noise is that it allowed us to focus on the mice’s voluntary (endogenous) attention, as opposed to involuntary (exogenous) attention recruited by the 3-bar grating.

We first extracted the spatial frequency and orientation energy by convolving Gabor filters with the black and white checker pattern in each stimulus frame (**Fig. 2A**). For example, our coherent grating produced maximum energy using a vertical Gabor with a spatial frequency of 0.077 cpd but produced little energy using a horizontal Gabor at the same spatial frequency (**Fig. 2A**). A typical FA lick is shown in **Fig. 2B** along with the preceding checkerboard frames and their vertical and horizontal energy that appeared on the cued side. In this example, increased vertical energy located in the nasal half of the stimulus happen to occur ∼0.33 seconds before the FA lick (**Fig. 2B, arrow**). By comparison, there was less horizontal energy in the stimulus before the same lick.

**Figure 2.**
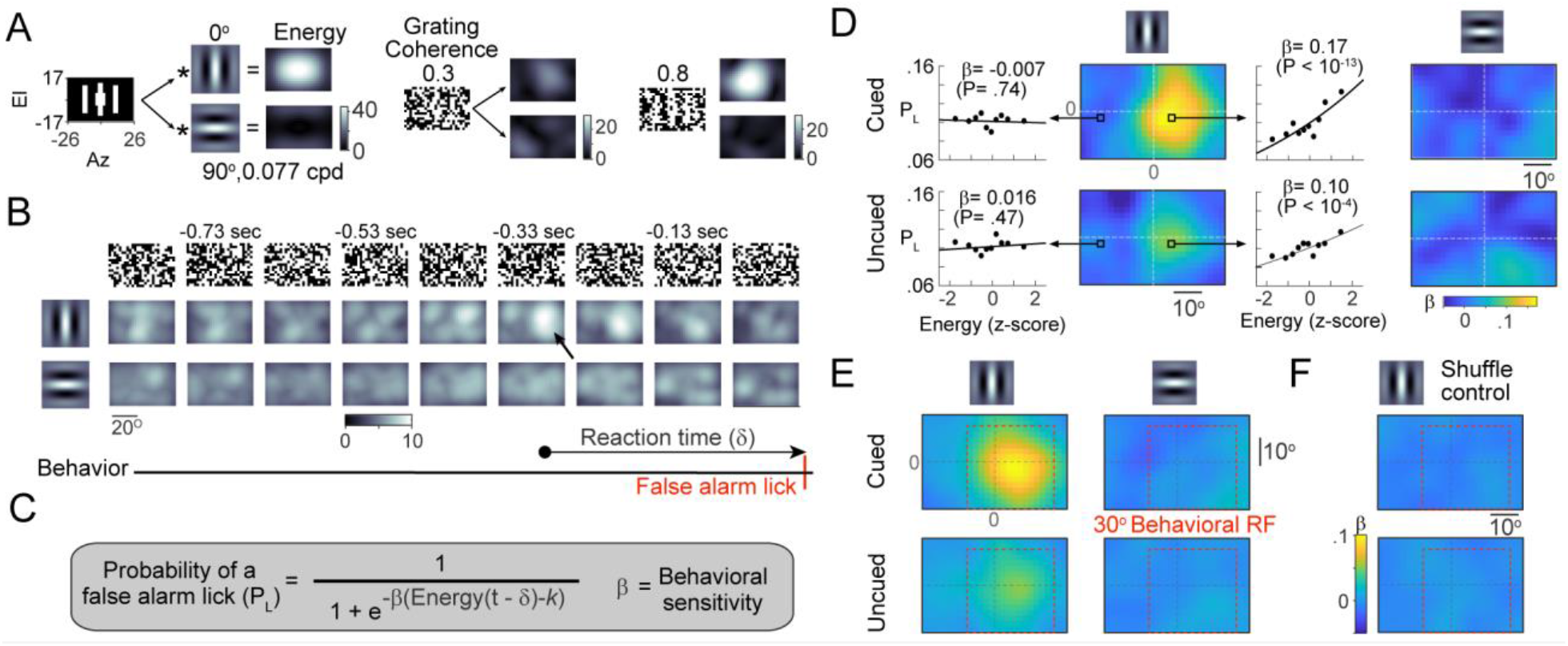
Behavioral reverse correlation of visual features and locations. **A**. Examples of convolving our 3-bar grating and checkerboard stimuli with either vertical or horizontal Gabors (0.077 cpd). **B-C**. Example checkerboard noise frames and energy presented on the cued screen before a false alarm (FA) lick. The probability of a FA lick (*P*_*L*_) was modeled as a logistic function of stimulus energy appearing δ secs before the lick (**C**). The behavioral sensitivity term (β) captures the correlation between the energy and the probability of a FA lick. **D**. Example behavioral sensitivity (β) for vertical and horizontal oriented energy from the mouse shown in Fig. 1. Insets relate orientation energy and the probability of a false alarm lick for the indicated checker location. Fits computed using the logistic function are shown along with the probability of a FA lick (filled circles) as a function of stimulus energy. Energy was always z-scored before fitting with the logistic equation. P-values are probability of β = 0. **E**. Average behavioral sensitivity from all mice (N = 13) for vertical or horizontal stimulus orientations. Red boxes indicate the behavioral receptive field (RF) used for subsequent analyses. **F**. Behavioral sensitivity maps computed from shuffled checkerboard sequences do not show a link between orientation energy and lick probability.

We applied a variant of the reverse-correlation approach^31,34^, that used a logistic generalized linear model to link FA licks to the energy contained in our random checkerboards (**Fig. 2C**). The logistic equation describes the probability of observing a FA lick as a function of stimulus energy. The slope parameter (β), captured the correlation between stimulus energy and the probability of a FA lick. We refer to β as *behavioral sensitivity*. The δ parameter is the time it took for the stimulus energy to produce a FA lick (i.e., reaction time), while *k* is the energy associated with a FA lick probability = 0.5.

### Behavioral sensitivity was strongest at nasal locations on the cued side

Behavioral sensitivity (β) represents the correlation of a visual feature linked to the decision to lick (see **Fig 1A**, green arrows represent stronger β). For example, FA licks produced at random with no regard for the stimulus would have a behavioral sensitivity = Behavioral sensitivity > 0 suggests that stimulus energy is positively correlated with FA licks. The larger the behavioral sensitivity, the stronger the correlation between a particular stimulus energy and FA licks.

We first estimated behavioral sensitivity as a function of spatial location by optimizing the logistic function for each checker location using Gabor filters to extract either vertical or horizontal stimulus energy. The effects of attention were measured by separately estimating behavioral sensitivity using stimuli from either the cued or uncued sides.

Example behavioral sensitivity as a function of spatial location and cue side for vertical and horizontal stimulus energy are shown as heatmaps (**Fig. 2D**). Behavioral sensitivity was strongest in a slightly nasal region of the checkerboard for vertical energy presented on the cued screen (**Fig. 2D, top left**). For the uncued screen (**Fig. 2D, bottom**) and horizontal energy (**Fig. 2D, right**) on either screen, behavioral sensitivity was reduced. The insets show example fits of the logistic equation from four locations; plotting binned lick probabilities as a function of normalized energy shows that the logistic function captured the probability of a lick.

Averaging across all mice revealed an unexpected result. Although vertical energy was spatially available across most of the checkerboard (**Fig. 2A, left**), behavioral sensitivity appeared in a nasal region of the cued screen (**Fig. 2E, top left**, P < 0.0001 that the average of β in the red box is > 0, single tailed t-test, N = 13). We do not know why the mice biased their FA licks to vertical stimulus energy in this nasal region but discuss the possibility that it could be related to specializations in the mouse visual system (*see discussion*).

We found no behavioral sensitivity to horizontal orientation on the cued screen (**Fig. 2E**., **top right**, P = 0.4), and reduced sensitivity to vertical energy on the uncued screen (**Fig. 2E**., **bottom left**, P < 0.001). As a control, we eliminated the correlation between stimulus energy and FA licks by shuffling checkerboards (**Fig. 2F**, P = 0.99 & 0.57, cued & uncued sides, respectively).

### Mice are most sensitive to features that match the 3-bar grating

We wanted to know how behavioral sensitivity was linked to different stimulus features such as spatial frequency, orientation, and local contrast. To accomplish this, we first collapsed our data across spatial location by focusing only on the checkboards with strong behavioral sensitivity – referred to as the *behavioral receptive field* (RF, red square in **Fig. 2E**). The behavioral RF encompassed ∼ 50% of the checkers and we summed the spatial frequency and orientation energy over this region for each stimulus frame (**Fig. 3A**).

**Figure 3.**
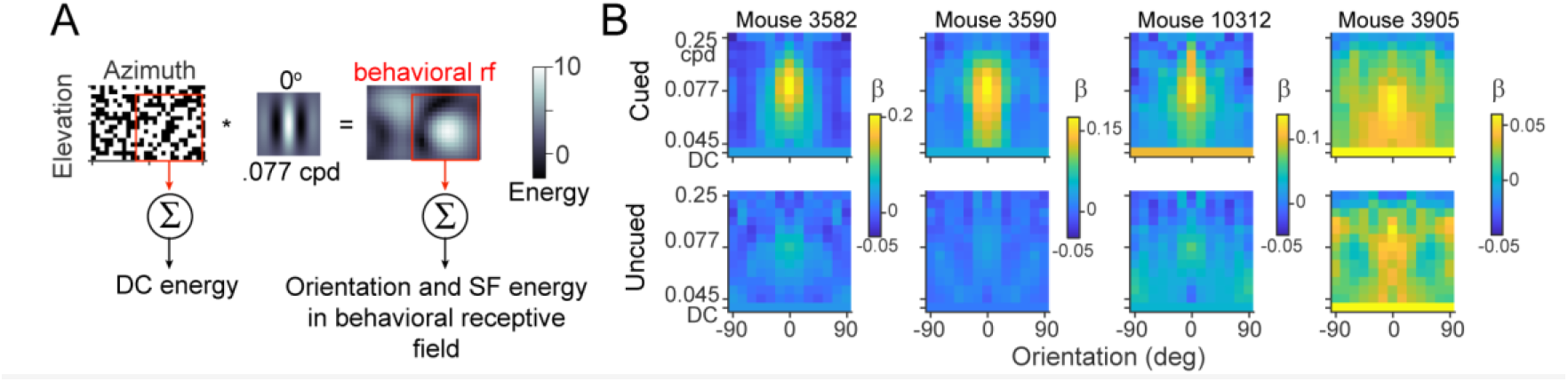
Behavioral sensitivity for orientation, spatial frequency and contrast. **A**. Flow chart showing the conversion of checkerboards into energy using Gabor filters of various orientation and spatial frequencies. Energy within the behavioral receptive field (red) was summed before estimating behavioral sensitivity. The number of white squares in the same region represents the local contrast and is referred to as DC energy. **B**. Representative behavioral sensitivity maps for 4 mice for a range of spatial frequencies and orientations on the cued (top) and uncued screens (bottom). Energy at 0 deg and 0.077 cpd corresponds to the peak oriented energy contained in our 3-bar grating.

The total number of white and black checkers was equal, regardless of the stimulus coherence. However, the 3-bar coherent grating produced local contrast in the number of white checkers that could been used by the mice during detection. Thus, we also computed the number of white checkers in the behavioral RF to capture the energy associated with the local contrast (referred to as the DC energy, **Fig. 3A**).

The behavioral sensitivity to spatial frequency, orientation, and DC energy of four well-trained mice is shown in **Figure 3B** (arranged in decreasing sensitivity, note the scale bar for each mouse). Animals generally had the highest behavioral sensitivity to orientation and spatial frequencies that corresponded to the peak energies in the 3-bar grating. Peak behavioral sensitivity to vertical energy was usually higher than it was to DC energy. In agreement with the effects of cueing shown above, the uncued screen had a reduced behavioral sensitivity profile in all but the animal with the weakest sensitivity (**Fig. 3B right**).

### Behavioral sensitivity develops over training and follows detection performance

Average behavioral sensitivity evolved over training (**Fig. 4A**). During the first 10 training sessions, FA licks showed no correlation with stimulus energy. Behavioral sensitivity to orientations and spatial frequencies contained in our 3-bar grating began to emerge during sessions 11 to 20 for both the cued and uncued screens. As training progressed, behavioral sensitivity increased mainly for stimulus energy located on the cued side. Sensitivity to the local contrast (DC energy) also emerged during training. It is worthwhile to note that the strength of our mouse behavioral sensitivity to orientation and spatial frequency is similar in magnitude to that reported for humans^30,31,35^.

**Figure 4.**
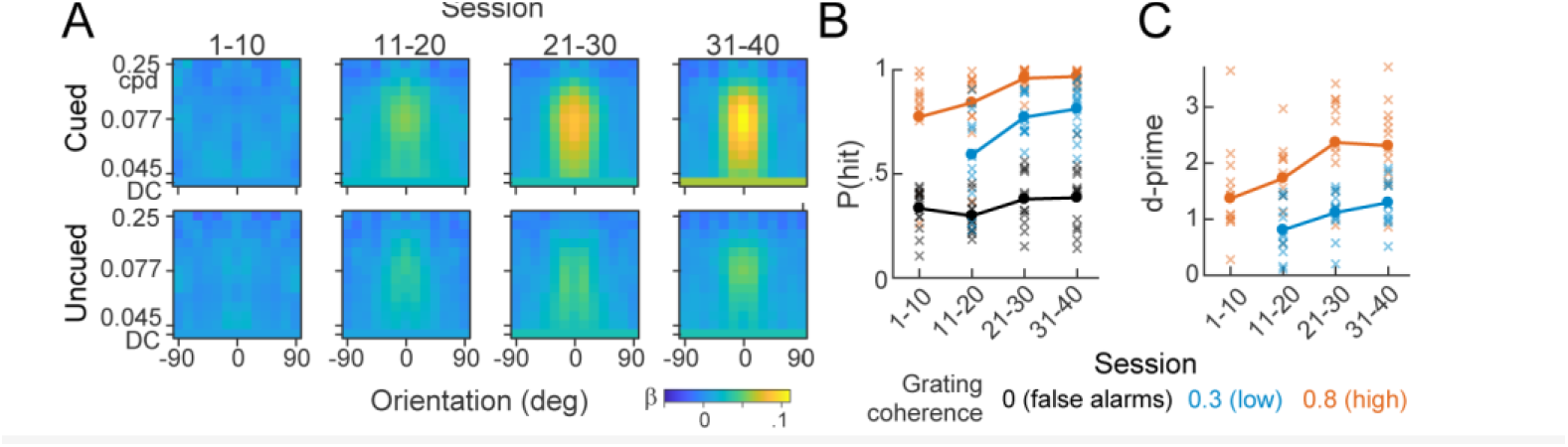
Behavioral sensitivity evolved during training and predicted detection performance. **A**. Average population behavioral sensitivity maps (N = 13) computed across the indicated sets of training sessions. Sensitivity to the vertically oriented 3-bar stimulus grows over training. **B**. Proportion of correct trials (hits) as a function of training sessions for a low (.3) and high (.8) grating coherence. False alarms are also shown (hits at grating coherence = 0). **C**. Behavioral d-prime computed from data in B. Behavioral performance grows over training sessions. Note that we only presented high grating coherences during the first few sessions to aid in early training.

The evolution of the population behavioral sensitivity suggests that training leads mice to preferentially monitor features that improved their ability to detect the 3-bar grating. To further examine this idea, we report detection performance (average hits and d-prime for valid cues, solid lines) for two grating coherence levels (low = 0.3 and high = 0.8) during the same training sessions (**Fig. 4B-C**). Detection performance improved over training, especially for the low coherence gratings), but false alarms (hits during a grating coherence = 0) stayed relatively constant. Thus, both behavioral sensitivity and grating detection improved for the cued side during training.

### Behavioral sensitivity is multiplicatively enhanced by attention

To reveal how behavioral sensitivity tuning to orientation and spatial frequency was affected by attention, we fit Gaussians to the population data in **Fig. 4B** sliced at 0 deg and 0.077 cpd **(Fig. 5A**, 2σ widths are shown). By the last training sessions, peak behavioral sensitivity to the cued side had increased to twice that of the uncued side (P = 0.0025, paired t-test). For these well-trained mice (sessions 31 to 40), the effect of cue location scaled the Gaussian fits without changing their width (orientation P = 0.46; spatial frequency P = 0.36; bootstrap). Thus, spatial attention directed at the cued side multiplicatively scaled behavioral sensitivity to orientation and spatial frequency by about a factor of two.

**Figure 5.**
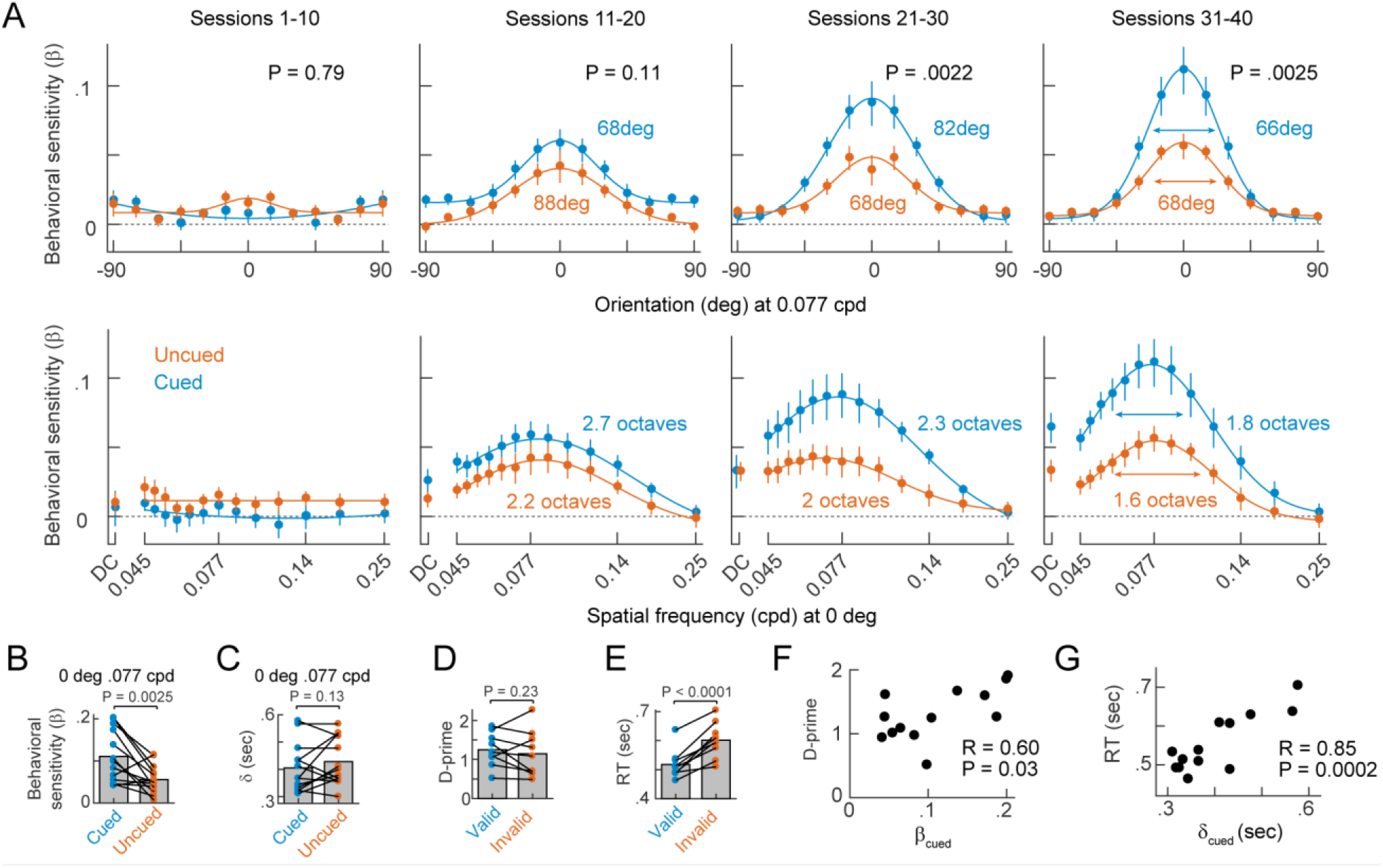
Attention enhances behavioral sensitivity. **A**. Population tuning curves of behavioral sensitivity versus stimulus orientation at 0.077 cycles per degree (top) and versus spatial frequency at an orientation of 0 degrees (bottom). Note spatial frequency is plotted on a log scale. The 2σ width of the fitted gaussians are labeled (except for the first training sessions that showed no tuning). **B-C**. Reverse-correlated measurements computed at 0 deg and 0.077 cpd show behavioral sensitivity (β) and reaction time (δ) for the cued and uncued sides. **D-E**. D-prime detection performance and reaction time (RT) for detection of the 3-bar grating (coherence = 0.3) for valid and invalidly cued trials. **F**. Behavioral sensitivity at 0 deg and 0.077 cpd on the cued side was correlated with d-prime detection of the validly cued 3-bar grating (coherence = 0.03). **G**. Reverse-correlated reaction time (δ) is correlated with the reaction time (RT) for detecting the validly cued 3-bar grating (coherence = 0.03). Points are individual animals in panels B-G. P-values are single-tailed paired t-test in panels A-E. R is Pearson’s correlation with associated P-value reported in panels F-G.

Furthermore, average behavioral sensitivity to local stimulus contrast (DC energy) was consistently less than sensitivity to energy at 0 deg and 0.077 cpd (**Fig 5A, bottom row**). This suggests that intensity was also a feature that the animals used to detect stimuli. This sensitivity also evolved as training progressed and was modulated by attention. But local contrast was not nearly as strongly linked to FA licks compared to oriented energy at 0 deg and 0077 cpd.

### Reverse correlation and invalid cueing measured different aspects of attention

The strong effect of spatial attention on population behavioral sensitivity was evident in individual mice, where all but two animals showed increased behavioral sensitivity at the cued side (**Fig 5B**). When estimating behavioral sensitivity (β), we also computed the time it took for stimulus energy to trigger a FA lick (δ in the logistic equation shown in **Fig 2C**), but this parameter was not modulated by attention (**Fig. 5C**). As a validation of our reverse correlation approach, we also included valid/invalid cueing on ∼10% of trials at the low grating coherence = 0.3 in nine animals (*see Methods*). Although these animals had no significant increase in d-prime detection between valid and invalid cues (P = 0.23) (**Fig. 5D**), they showed a consistent increase in their reaction time to the 3-bar grating when the cue was invalid (P < 0.0001) (**Fig. 5E**).

Both reverse-correlation and valid/invalid cueing revealed robust effects of attention on different aspects of the mice behavior: Changes in sensitivity were found with reverse correlation (**Fig 5B**), whereas changes in reaction time were found with invalid cueing (**Fig 5E**). We speculate that this difference is because gratings occurring on the uncued side (invalidly cued trials) exogenously captured the animals’ attention. When using reverse correlation, the checkerboard energies on the uncued screen were much too weak to exogenously capture attention (see Discussion).

Finally, we wanted to know if our two measurements from the reverse correlation, behavioral sensitivity (β) and reaction time (δ), were related to the detection of the 3-bar grating. While behavioral sensitivity at 0 deg and 0.077 on the cue side was moderately predictive of detection of the low-coherence 3-bar grating (**Fig 5F**), our reverse-correlated reaction time, was strongly correlated with the time it took for mice to lick in response to the 3-bar grating (**Fig 5G**). Thus, mice that were slow/fast to FA lick in response to checkerboard noise were also slow/fast to lick in response to the 3-bar grating. Taking these results together, our reverse-correlated measurements (β and δ) reflected detection capability, endogenous attentional modulation, and the speed of behavioral responses in our mice.

### Mouse behavioral sensitivity approached that of an ideal observer

Although it makes sense that peak behavioral sensitivity (β) corresponded to the orientation and spatial frequency of the peak energy contained in our 3-bar grating, several questions remain. Why did mice have behavioral sensitivity at other orientations and spatial frequencies? And why did mice show a behavioral sensitivity to the local stimulus contrast (DC energy) that was about half as strong as their peak sensitivity? To answer these questions, we used an ideal observer model.

Briefly, our ideal observer model used signal detection theory to discriminate between random and 0.3 coherence checkerboards using the same Gabor filters as the reverse-correlation analysis. At each orientation and spatial frequency, the ideal observer summed the energy across the same behavioral RF as that of the mice (**Fig. 6A**), and a d-prime sensitivity was computed based on the energy distributions (**Fig. 6B**) derived from 50,000 stimulus frames. The resulting d-prime map in **Fig. 6C** illustrates how well the ideal observer discriminated 3-bar checkerboards using different features.

**Figure 6.**
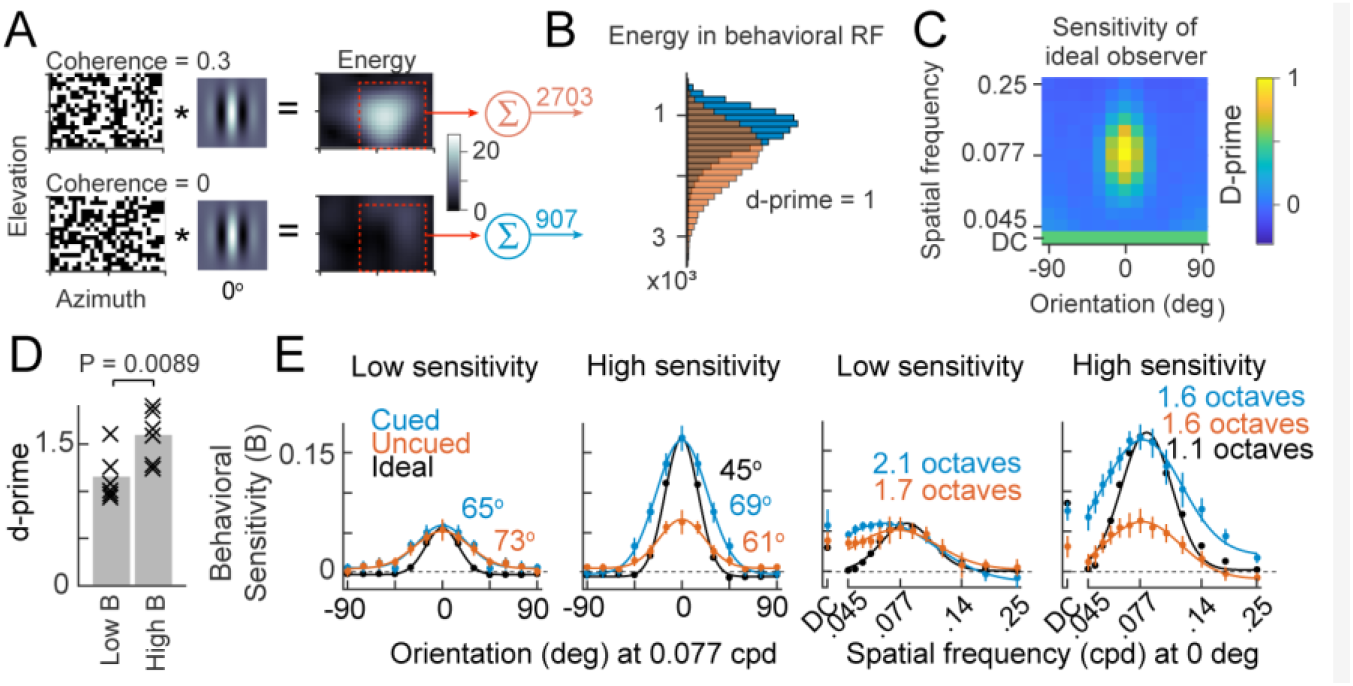
Ideal Observer analysis. **A-B**. Example of the ideal analysis at a single orientation and spatial frequency. Checkerboards with the 3-bar grating (coherence = 0.3) or those with pure noise were convolved with a vertically oriented Gabor (0.077 cycles per degree). Red boxes indicate the behavioral RF from which orientation energy is summed to create two histograms in panel B whose spacing corresponded to a d-prime of 1 (see Methods). d-prime corresponds to the performance of an ideal observer to discriminate the noise and grating checkerboards based on random draws from each distribution. **C**. Heatmap shows d-primes computed from using Gabor filters with the indicated spatial frequencies and orientations. DC sensitivity is based on the number of white squares in the behavioral RF. **D**. Detection of 0.3 coherence gratings for mice with low (β < 0.1) and high (β > 0.1) behavioral sensitivity on training sessions 31-40. **E**. Tuning curves of the ideal observer sensitivity (d-prime) and behavioral sensitivity (β) for the mice in panel D versus grating orientation, spatial frequency and DC. Note that the ideal observer tuning was scaled to match the mice peak cued-side behavioral sensitivity and was taken from slices of the data shown in C.

As expected, energy at 0 deg and 0.077 cpd contained the most information about the presence of the 3-bar grating. Nearby orientations and spatial frequencies were also informative (d-prime > 0), likely because our Gabors are not perfect bandpass filters. The ideal observer model also reported sensitivity at DC that was about half that of the peak orientated sensitivity at 0.077 cpd.

We wanted to compare the ideal observer to the “most sensitive” mice (largest β in **Fig 5B**) because we imagined these animals were the most optimal in how they used stimulus features^36^. To examine this, we evenly divided animals by their behavioral sensitivity at 0 deg and 0.077 cpd (**Fig 6D**) and compared the tuning of these groups to that of the ideal observer. This comparison revealed several interesting features. First, and as shown above in Fig. 5F, high-sensitivity mice tended to also be good detectors of the 3-bar grating (**Fig. 6D**). Second, high-sensitivity mice qualitatively chose the same strategy as the ideal observer and monitored orientations and spatial frequencies contained in the 3-bar stimulus (**Fig. 6E, right**). Third, high-sensitivity mice monitored the local stimulus contrast (DC energy) to the same extent as predicted by the idea observer. Fourth, high-sensitivity mice showed strong attentional effects by focusing much more on the cued side checkerboards (which an ideal observer would also do). Thus, high sensitivity mice resembled an ideal observer, but with wider tuning. Low sensitivity mice were less like the ideal observer. These animals emphasized orientation, spatial frequency, and stimulus contrast (DC energy) equally (**Fig 6E, left**) and showed no appreciable attentional effects.

Taken together, these data show that our high-sensitivity mice found oriented stimulus energy more informative than stimulus contrast for detecting the 3-bar grating. These animals also better understood the saliency of the cue.

## Discussion

We demonstrated that behavioral reverse-correlation^7,28–33,35,37^ measures the spatial- and feature-based properties of attention in mice. We trained mice to detect three vertical bars in a background of random checkerboard noise. Mice saw such noise on two screens but were cued to which screen would show the three bars. Mice were highly motivated to detect the stimuli which led them to mistakenly ‘see’ the weak bars in the checkerboard noise. We leveraged this false alarm behavior to show that mice attended to visual features that contained evidence of the 3-bar stimulus. To our knowledge, this is the first demonstration that mice can simultaneously attend to different spatial locations and features of a visual stimulus.

The resulting maps of behavioral sensitivity showed that mice attended a small patch of the cued checkerboards for vertically oriented energy whose spatial frequency matched that of our three vertical bars. This patch represented the attentional spotlight and we discovered that it evolved with training and followed the audiovisual cue. An ideal observer performing the same task revealed sensitivity to visual features that were like those found in mice but were more narrowly tuned.

Multiplicative interactions are well-known to describe how neurons in visual cortex represent multiple features^38–41^. Such interactions are a major proposed mechanism for the attentional enhancement of neural responses^2,2,3,6,42^ and behavior^7,8,28,35,37,43,44^. Our reverse correlation method allowed us to test many stimulus features and locations at once and revealed how mice weigh these variables to make behavioral decisions. Our study is the first to measure these weights in mice, which were surprisingly similar to that found in humans^37^. These results suggest that multiplicative models of attention developed in humans and primates are likely to apply in mice.

### Comparisons with other rodent attentional tasks

There are now a few tasks to measure attention in mice^16–24^. Several use implicit methods to direct the locus of spatial attention^16,23,45^. For example, stimuli that appear in one location for blocks of trials before unexpectedly switching to a new location lead mice to detect poorly at block transitions because their attention is misallocated^16^. As another example, stimuli that appear more often in one of two locations lead some mice to monitor only the high likelihood location but lead others to switch between high and low likelihood locations^23^. These studies test the effects of spatial attention on a single stimulus feature and do not reveal tuning.

A small set of studies have used explicit cues to direct a mouse’s attention and observe its underlying neural correlates^17,19–21^. This classical design allowed them to make the spatial cue either valid or invalid and confirm that mice, like primates, experience increased stimulus detection and faster reaction times with cued attention^19^. Attention in these studies improved mouse performance by ∼30%. Our own invalid cueing produced comparable effects on reaction times, however, our reverse correlation method found attention scaled the sensitivity to features more than 100%. We suspect the reason for this difference is because attentional effects on behavior are strongest when using weak stimuli.

### Endogenous vs exogenous attention

Strong stimuli in uncued locations are well known to transiently take attention away from the cued side^3,46,47^. Our checkerboards were unlikely to be strong enough to capture attention in this way, and thus our cue effects on behavioral sensitivity are consistent with sustained voluntary attention (endogenous attention). By comparison, the valid/invalid cueing effects we observed could have been due to involuntary attention (exogenous attention). This is because the appearance of the 3-bar stimulus on the uncued side was probably strong enough to capture attention. Reorienting of attention takes time, and we observed consistent increases in reaction times when mice responded to the 3-bar stimulus on the uncued side during invalidly cued trials.

### Of mice and ideal observers

Why was the behavioral sensitive tuning of mice more broadly tuned than the ideal observer? Both the mice behavioral sensitivity (β) and the ideal observer sensitivity (d-prime) were based on the same Gabor filters. We suspect the difference lies in the relatively broad tuning of mouse visual cortical neurons^48–50^. A quick survey of recently published tuning widths shows that the orientation tuning width of mouse V1 neurons is ∼40-50 degrees^48,51^ while spatial frequency tuning widths are ∼2-3 octaves^49,52^. In addition, neurons in the mouse visual system respond to much lower spatial frequencies compared to humans (.01-.08 cpd^49^ versus 0.5-5 cpd^53^, respectively). Although our mouse reverse-correlated behavioral sensitivity to orientation and spatial frequency had similar behavioral sensitivity magnitudes compared to humans, the spatial frequency range reported and orientation tuning width for humans was much higher^30,31,35,37^.

Only a few studies have addressed how a visual cortical neuron’s link to behavior changes as a function of its stimulus tuning. For example, Bosking and Maunsell demonstrated that the correlation between a directional selective visual neuron’s activity and behavior gradually fell off as the motion stimulus direction moved away from that preferred by the neuron^54^. This suggests that downstream areas involved in generating behavioral responses ‘weigh’ a visual neurons activity proportional to how much information the neuron provides about the stimulus. Thus, the wider behavioral sensitivity tuning of mice compared to the ideal observer could be explained by the fact that mouse visual neurons are more widely tuned than the Gabor filters used by the ideal observer model. Combining our method with in vivo calcium imaging of mice V1 could allow a direct test of this idea.

### Behavioral sensitivity and attention in mice and primates

We specifically chose a stimulus with narrowband orientation and spatial frequency energy similar to that used to measure attention in humans^28,30,31,35,37^. We note that the behavioral sensitivities and effects of attention we see in mice are remarkably similar to those obtained in human^30^. While there are substantial differences in the retinal specializations of humans and mice^55^, we did notice high behavioral sensitivities located in a slightly anterior location of our two monitors. This region abuts the binocular zone of mice^55^, which in recent years, is receiving increasing attention as a functional fovea^56–59^. Retinal ganglion cells types innervating the primary visual thalamus are particularly enriched in this region^57,60^ and it is neurally magnified in mouse visual cortex^56^. Moreover, other studies employing freely moving prey capture behaviors suggest that mice actively stabilize this part of their retina and require it for the final stages of prey capture^58,59^. Our results showing an attentional focus in a very similar region could reflect a strategy to use this retinal specialization to pay attention. Such a strategy is common in humans where eye-movements are a well-known measure of attention^1,30,45^. Future work could use our attentional task and reverse-correlation methods together with eye-tracking to address this idea.

## Methods

### Animals

Animals were used in accordance with the rules and regulations established by the Canadian Council on Animal Care and protocols were approved by the Animal Care Committee at McGill University. Male and female C57BL6/J mice aged 60-120days were implanted deeply anesthetized using isofluorane (5% induction, 1-3% maintenance), mounted on a stereotaxic frame (Stoelting), and eye ointment was applied to prevent corneal drying. A circular incision was made along the scalp using micro scissors to remove the scalp. Lidocaine was applied to skull surface before the underlying fascia was cleaned and the skull was dried. Bregma and lambda were then levelled to within 0.1mm of one another. A custom designed stainless steel headplate was cemented directly to the skull using Metabond (C&B). Following surgery, the scalp was sutured closed, the animal removed from the stereotaxic frame and placed on a heat pad for recovery, and analgesia administered subcutaneously for 3 days following surgery.

### Behavioral Arena

The behavioral apparatus consisted of a custom built soundproofed dark box that displayed visual stimuli via two 60 Hz LCD displays angled at 32 degrees from the mouse’s midline. The head fixed mouse was placed in the center of the apparatus on a custom designed, 3D-printed, platform. A lick spout was positioned in front of the mouse’s mouth to administer the liquid reward. Licking was capacitively sensed using custom electronics and an Arduino Uno. Liquid reward was delivered by controlling a solenoid pinch valve. The behavioral task and presentation of visual stimuli was controlled using the psychophysics toolbox extensions for Matlab (The Mathworks), and custom data acquisition software.

### Training schedule

Mice were housed in a 12hr/12hr reversed light-dark cycle, with lights turned off in the morning. After recovery from the head plating surgery, mice were initially head-fixed in behavioral rigs to habituate to the setup. In these sessions (up to 3), free liquid rewards were given to the mice to associate reward with the lick spout. Once training on the grating stimulus began, home-cage water bottle was removed and a minimum daily quota of 1mL water was enforced. Animals either drank this amount during training or were completed to this amount, if necessary, at the end of the day. A typical training session lasted 1-2 hours or until the mouse ceased self-initiating trials. Animals were weighed daily, and the amount of water consumed during each training session recorded (typically .7-1.2mL). We gradually increased the wait times and grating coherence (Fig. 1E) while monitoring the animals detection performance. By training session 10, mice reliably produced psychometric curves. As shown in **Fig. 5B-C**, detection performance, especially at the low coherence level of 0.3, continued to improve over the 40 training sessions used in this study. Mice were never rewarded for false alarm licks, or licks in response to a zero coherence grating.

### Visual stimulus

Dynamic checkerboards were updated at 30 Hz. Each checkerboard was 17 (elevation) X 26 (azimuth) checkers and individual square checkers were 2° X 2° (**Fig. 1C-D**). Positive azimuth corresponds to the nasal direction. A visual cue was used to guide the animal’s attention to one monitor and consisted of a black and white frame that surrounded the checkerboard and a small black and white rectangle placed on one side of the checkerboard (**Fig. 1E**). Simultaneously, an audio cue was also presented from a speaker mounted behind the monitor and was an 8Khz tone at ∼75dB SPL. Once a start lick occurred, both the visual and audio cue simultaneously faded linearly over 6 secs.

The coherent stimulus consisted of three vertical white bars combined with a noisy background (**Fig.1C-D**). The 3-bar stimulus was derived from a thresholded vertical Gabor (.077 cpd). For random noise (i.e., coherence = 0), all checkers were randomly assigned to black or white on each frame. The coherent grating stimulus was generated using three steps: 1) The three vertical bars in the checkerboard were set to white, and all other checkers set to black (see grating in **Fig. 1C**). 2) Black checkers were then randomly flipped to white so that the number of white and black checkers were equal. This corresponded to a grating coherence = 1 (**Fig. 1D**, far right). 3) For grating coherence < 1, a number of checkers, proportional to one minus the coherence, were randomly assigned either black or white (referred to as noise checker). Thus, for a grating coherence = 0.5, half the checkers are noise (**Fig. 1D**). This checkerboard stimulus maintained, on average, an equal number of white and black checkers for all grating coherence.

### Trial design and cueing

Mice performed a spatially cued detection task (**Fig. 1E**). They were trained to lick in response to the onset of the 3-bar coherent grating stimulus. Trials began with the onset of the audiovisual cues and a static random checkerboard; cues and coherent stimuli changed sides together every ∼25 completed trials (i.e., trials that ended in either a hit or miss). Mice were required to produce one or more start licks within 10 sec. If no start lick occurred, the trial was scored as an *ignore*. After the start lick, dynamic random checkerboard noise began on both screens. The coherent grating then occurred on the cued screen at a random time. The minimum and maximum wait times for the coherent grating to occur was gradually lengthened during the initial training sessions until it reached 4 – 12 secs (∼session 15). The time the coherent grating appeared was randomly chosen between the minimum and maximum wait times (based on an exponential distribution in order to produce a flat hazard function^61^).

Once the coherent grating occurred, a lick that occurred within a 0.2 to 1.5 sec reaction time window was scored as a *hit* and a reward was provided after a 0.3 sec delay. Licks that occurred before 0.2 sec were scored as a *false alarm* and the trial ended with no reward. No licks, or licks occurring after 1.5 sec, were scored as *miss* and no reward was provided.

When mice licked before the onset of the 3-bar coherent grating it was scored as a false alarm (*FA lick*) and the trial restarted. Restarts cause a static checkerboard to appear on the cued side along with the audiovisual cues for 0.5 sec (the uncued screen was gray), followed by dynamic checkerboard noise. Trials were allowed to restart for four minutes, after which the trial ended with no reward. Mice rarely produced enough FA licks to extend the trial beyond 20 sec.

During the first 20 training sessions the coherent grating always appeared in the cued checkerboards. In 9 of 13 mice, an invalid cueing paradigm was used as an additional measure of spatial attention during training sessions 20 to 40. For invalidly cued trials the 3-bar coherent grating (0.3 coherence) occurred on the uncued side and were ∼10% of all trials (i.e., the cue was valid on 90% of trails).

### Reverse correlation

We linked FA licks to stimulus features and locations using reverse correlation^34,37^. In this variant of the reverse correlation approach, FA licks are modeled using the logistic function shown in **Fig. 2C**. In this equation, the probability of a FA lick (*P*_*L*_) is a function of a stimulus feature (referred to as *Energy* in our logistic equation). In our study, stimulus energy was characterized as orientation, spatial frequency and local contrast. The three unknown parameters of the logistic equation were: β the behavioral sensitivity of FA licks to the stimulus energy, δ the time it took stimulus energy to produce a FA lick, and *k* which defines the stimulus energy corresponding to *P*_*L*_ = 0.5. β was our most important parameter as it captured the correlation between a stimulus feature and behavior. A β = 0 indicated that stimulus energy had no correlation with FA licks. A β > 0 indicated that increased stimulus energy was positively correlated with the probability of a lick, while β < 0 suggested increased energy was associated with a reduction in the lick probability.

On each random checkerboard frame, stimulus energy was computed for orientation, spatial frequency and local contrast. We used quadrature phased Gabor filters convolved with each stimulus frame to extract energy associated with orientation and spatial frequency^35^. These Gabor filters were 26 × 26 checkers in size constructed with a standard deviation = 4 checkers (8 deg). We used 12 orientations from 0 (vertical) to 165 deg in 15 deg increments, and 13 spatial frequencies from 0.046 to 0.25 cpd with variable increments. Because orientation is symmetric with respect to vertical, we combined like-orientation energies before solving for the parameters in the logistic equation (e.g., 15 and 165 deg, 30 and 150 deg, etc.)^35,37^. Combining like-orientations increased the number of data points and thus improved the accuracy of behavioral sensitivity estimates and did not qualitatively change our results.

Convolving Gabors with our checkerboards produced orientation and spatial frequency energy centered at each checker (**Fig. 2A-B**). Thus, our first reverse correlation analysis extracted behavioral sensitivity (β) across stimulus space (**Fig. 2D-F**). Subsequent analyses collapsed across space by summing energy within a behavioral RF defined by the red box in **Fig. 2E-F** before solving the logistic equation (**Fig. 3A**).

Although our checkerboard stimulus maintained the same number of black and white checkers regardless of grating coherence, local differences in the number of white checkers (contrast) occurred for checkerboards with grating coherences > 0. Mice could have been detecting these contrast changes during FA licks. Thus, we also used the number of white squares in the behavioral RF as a feature of our stimulus (referred to as DC energy, **Fig. 3A**).

We aligned our stimulus energy to FA licks. We only included FA licks that were preceded by at least 1.5 sec of random checkerboard noise. The average duration of checkerboard noise before a FA lick was 3.8 ± 0.7 sec (SD, N = 13). For each FA lick we temporally filtered the stimulus energy with a centered Gaussian filter (σ = 3 stimulus frames). We then resampled the stimulus energy every 10 frames starting at the FA lick minus the reaction time (δ). The temporal filtering and resampling greatly reduced the noise in our estimate of behavioral sensitivity and represented the temporal integration in the mouse visual system.

For a given reaction time (δ), a generalized linear model (GLM) approach was used to solve for β and *k* in our logistic equation^35,37^. We first divided our FA licks into three data partitions by assigning every 3^rd^ lick to a different partition. After temporally filtering and resampling the data, we then estimated the parameters of the logistic equation using a three-step process: First, from one data partition, solve the GLM model for every reaction time (δ), where δ ranged from 8 to 20 frames (0.267 to 0.667 sec). Select the highest β as the reaction time δ_i_ to use in step 2. Second, using the reaction time (δ_i_) from step 1, solve the GLM model for β_i_ using the data in the other two partitions. And third, repeat these two steps until all possible combinations of the data partitions have been used to estimate δ_i_ and β_i_.

Our reported estimates for behavioral sensitivity (β) and reaction time (δ) were the averages of δ_i_ and β_i_. Partitioning our data ensures that our estimates of behavioral sensitivity were unbiased. In other words, this method ensured that if mice randomly FA licked with no regards to stimulus energy, then behavioral sensitivity would approach zero. This was verified in our shuffle control (**Fig. 2F**).

### Ideal observer model

This model computed the sensitivity (d-prime) of an ideal observer that discriminated random checkerboards (coherence = 0) from checkerboards containing the weak 3-bar grating (coherence = 0.3). The ideal observer used the same stimulus features as the mouse reverse-correlation analysis (orientation, spatial frequency and DC energy located in the behavioral RF, **Fig. 3A**). For each stimulus feature, two energy distributions were computed using 50,000 stimulus frames (**Fig. 6A-B**). The d-prime sensitivity of the ideal observer was computed using the standard expression 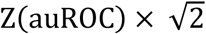
, where auROC is the area under the receiver operating characteristics curve for the two distributions and Z is the inverse of the standard normal cumulative distribution function^62^.

